# Morphometry of SARS-CoV and SARS-CoV-2 particles in ultrathin plastic sections of infected Vero cell cultures

**DOI:** 10.1101/2020.08.20.259531

**Authors:** Michael Laue, Anne Kauter, Tobias Hoffmann, Lars Möller, Janine Michel, Andreas Nitsche

## Abstract

SARS-CoV-2 is the causative of the COVID-19 disease, which has spread pandemically around the globe within a few months. It is therefore necessary to collect fundamental information about the disease, its epidemiology and treatment, as well as about the virus itself. While the virus has been identified rapidly, detailed ultrastructural analysis of virus cell biology and architecture is still in its infancy. We therefore studied the virus morphology and morphometry of SARS-CoV-2 in comparison to SARS-CoV as it appears in Vero cell cultures by using conventional thin section electron microscopy and electron tomography. Both virus isolates, SARS-CoV Frankfurt 1 and SARS-CoV-2 Italy-INMI1, were virtually identical at the ultrastructural level and revealed a very similar particle size distribution with a median of about 100 nm without spikes. Maximal spike length of both viruses was 23 nm. The number of spikes per virus particle was about 30% higher in the SARS-CoV than in the SARS-CoV-2 isolate. This result complements a previous qualitative finding, which was related to a lower productivity of SARS-CoV-2 in cell culture in comparison to SARS-CoV.

## Introduction

The Severe Acute Respiratory Syndrome Coronavirus 2 (SARS-CoV-2) is a *Betacoronavirus* which entered the human population most probably at the end of 2019 and is spreading pandemically around the world^1^. The virus causes the disease termed COVID-19 which primarily affects the respiratory system^1,2^ but can extend to other organs^3^. Severity of the disease is highly variable from non-symptomatic to fatal outcomes^1^.

SARS-CoV-2 is genetically similar to SARS-CoV (79% sequence identity^4^) which appeared in the human population in 2003. Both viruses use the same receptor (i.e. the angiotensin-converting enzyme 2, ACE2) for host cell entry^5^. Infection of different cell lines and of patient material could be shown^6,7,8^. Ultrastructural hallmarks of entry, replication and assembly seem to be virtually identical to SARS-CoV^9^. Like all viruses of the family *Coronaviridae*, the virus is a biomembrane-enveloped virus with prominent surface projections, called spikes or peplomers, which are formed by a glycoprotein (S protein) trimer (Fig. 1). The molecular structure of the spike protein was already resolved by cryo-electron microscopy (EM)^10^. The virus genome is a single plus-strand RNA molecule which is associated with the nucleoprotein (N protein) in the enveloped lumen of the virus (Fig. 1).

**Figure 1.**
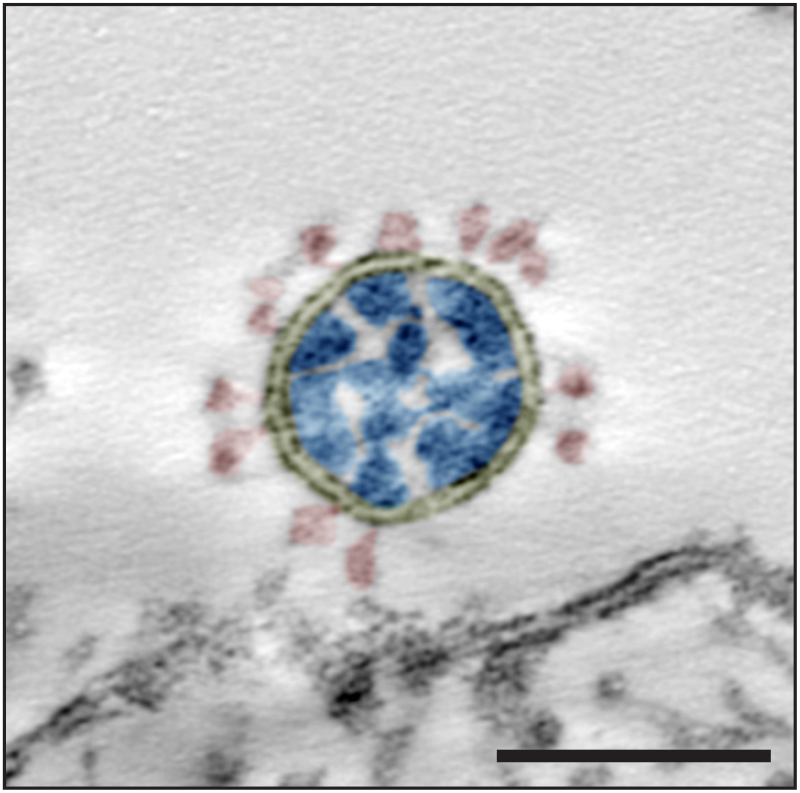
Transmission EM of a single virus particle of SARS-CoV-2 at the surface of a Vero cell in an ultrathin plastic section (10 summed up digital slices of an electron tomogram). The section through the virus particle shows the main ultrastructural features of the virus which were manually highlighted by color: yellow = virus-enveloping membrane, red = surface projection (spike, peplomer; trimeric S protein), blue = ribonucleoprotein (N protein and RNA). Scale bar = 100 nm.

Very recently, morphometric data on isolated SARS-CoV-2 particles^11–13^ and virus particles in cells^14^, obtained by cryo-EM, were published or became available as a preprint. While cryo-EM is definitely the best method to study virus ultrastructure and structural biology, conventional EM, using plastic embedding, still is of relevance, especially for the study of samples, which cannot be easily analyzed by cryo-EM, such as complex multicellular objects or pathological material obtained from patients. Search of viruses in such material is difficult and needs a suitable reference obtained with virus infected cell culture material using the same preparation technique^15^. To provide reference data for this purpose, we carried out a study on the morphometry of virus particles of SARS-CoV-2 in comparison to virus particles of SARS-CoV by using transmission EM of the virus in thin sections of plastic embedded infected cell cultures.

We show the particle size distribution of virus particle profiles in conventional ultrathin sections and in single-axis tomograms of thicker sections. The spike number was determined for virus particle profiles in ultrathin sections and compared with measurements of the spike number of complete particles in tomograms. The study provides robust data, including all raw data files, on the morphometry of the two coronaviruses as they appear in conventional thin section EM of virus producing cell cultures and demonstrate that the investigated SARS-CoV and SARS-CoV-2 isolates are very similar in their ultrastructure apart from a small difference in their spike number.

## Methods

### Virus isolates

The following virus isolates were used:

1. SARS Coronavirus Frankfurt 1 (SARS-CoV)^16^
2. SARS Coronavirus 2 Italy-INMI1 (SARS-CoV-2)^17^

### Cell culture

Vero E6 cells (African green monkey kidney epithelial cell, ECACC, ID: 85020206) were cultivated in cell culture flasks with D-MEM, including 1% L-glutamine and 10% fetal bovine serum, for 1 d at 37°C and 5% CO_2_ to reach approximately 70% confluence. To infect the cultures with virus, the medium was removed and 10 ml of fresh medium with diluted virus stock suspension was added to the cells (the multiplicity of infection was about 0.01). After incubation for 30 min, as indicated above, 20 ml of medium was added and cells were further incubated. Cultivation was stopped 24 h after addition of the virus suspension by replacing the medium with 2.5% glutaraldehyde in 0.05 M Hepes buffer (pH 7.2). Incubation with the fixative lasted at least 1 h at room temperature. Fixed cells were scraped from the culture flasks and collected in centrifuge tubes.

### Electron microscopy (EM)

Fixed cells were sedimented by centrifugation (3000 g, 10 min) using a swing-out rotor and washed twice with 0.05 M Hepes buffer. The cell pellet was heated to 40 °C in a water bath and mixed with 3% low-melting point agarose (1:1 [v/v]) at 40 °C. After a brief (approx. 2-3 min) incubation at 40 °C, the suspension was centrifuged in a desktop centrifuge using a fixed-angle rotor for 5 min at 5000 g and cooled on ice to form a gel. The cell pellet was cut off from the agarose gel block by using a razor blade and stored in 2.5% glutaraldehyde in 0.05 M Hepes buffer. Postfixation, *en bloc* contrasting, dehydration and embedding in epoxy resin (Epon^18^) were done following a standard protocol^19^ (Supplementary Table 1).

Ultrathin sections were produced with an ultramicrotome (UC7, Leica Microsystems, Germany) using a diamond knife (45°, Diatome, Switzerland). Sections were collected on bare copper grids (300 mesh, hexagonal mesh form), contrasted with 2% uranyl acetate and 0.1% lead citrate and coated with a thin (2-3 nm) layer of carbon. For electron tomography, gold colloid (10-15 nm; 1:10 or 1:20 diluted) was added to the carbon-side of the sections by incubating the sections on a drop of the gold colloid suspension for 1-5 min at room temperature.

EM of thin sections was performed with a transmission electron microscope (Tecnai Spirit, Thermo Fisher Scientific) which was equipped with a LaB_6_ filament and operated at 120 kV. Magnification calibration of the microscope was done by using the MAG*I*CAL calibration reference standard for transmission EM (Technoorg Linda, Hungary). Images were recorded with a side-mounted CCD camera (Megaview III, EMSIS, Germany) and 1376 x 1032 pixel. Tilt series for electron tomography were acquired by using the tomography acquisition software of the Tecnai (Xplore 3D v2.4.2, Thermo Fisher Scientific) and a bottom-mounted CCD camera (Eagle 4k, Thermo Fisher Scientific) at 2048 x 2048 pixel. A continuous tilt scheme at one degree interval was used and at least 120 images were recorded (minimum +60 to −60°). Tracking before image acquisition was performed to compensate image shifts introduced by the mechanics of the stage. Alignment and reconstruction were done with the Inspect3D software (Version 3.0; Thermo Fisher Scientific) by using a defined procedure and the “[Simultaneous Iterative Reconstruction Technique](https://tomroelandts.com/articles/the-sirt-algorithm)” (SIRT) with 25 iterations (Supplementary Table 2).

Additional single-axis tilt series (at least −60° to 60°, increment 1°, defocus −0.2 μm) of thicker sections (200-250 nm) were acquired with a transmission electron microscope (JEM-2100, Jeol) at 200 kV with a pixel size of 0.57 nm by using a side-mounted CCD camera (2048 x 2048 pixel, Veleta, EMSIS, Germany) and SerialEM^20^ (version 3.7.11). Alignment of the tilt series (using 10 nm colloidal gold fiducials) and reconstruction of the tomograms were performed with the IMOD software package^21^ (version 4.9.12) using SIRT with 25 iterations after low pass filtering (cut off = 0.35, low pass radius sigma = 0.05) of the aligned image stack.

### Measurement of virus particle size

Size of virus particle profiles was measured in images of ultrathin sections (65, 85, 110 nm) and in tomograms of thicker (150-180 and 200-250 nm) plastic sections.

Extracellular virus particles in ultrathin sections were selected randomly at the microscope and recorded with the side-mounted camera (at a magnification of 105,000x), if they met the following criteria: (1) the particle was morphologically intact; (2) the particle was not pressed against other structures. Six datasets were recorded (see Table 1). Data set 1 and 2 were also used to measure the maximal length of the spikes (see below). For this purpose, virus particles were selected which, in addition to the two criteria mentioned above, are covered with spikes by at least 2/3 of their particle perimeter. Particle size measurements were done with Fiji^22^ by selecting the outer leaflet of the virus membrane with the “polygonal selection” tool and the measurement setting „fit ellipse“. Maximal and minimal diameter of the fitting ellipse and shape descriptors, such as aspect ratio and circularity (4π*area/perimeter^2^), were determined.

**Table 1.** Overview of the datasets used for virus particle measurements

| Data set # | Virus isolate | Sample | Number of Sections | section thickness [nm] | number of files | file format | pixel size [nm] |
| --- | --- | --- | --- | --- | --- | --- | --- |
| 1 | SARS-CoV Frankfurt | A | 4 | 65 | 126 | tif, 16 bit | 0.64 |
| 2 | SARS-CoV-2 INMI-Italy | B | 4 | 65 | 128 | tif, 16 bit | 0.64 |
| 3 | SARS-CoV-2 INMI-Italy | C | 5 | 65 | 122 | tif, 16 bit | 0.64 |
| 4 | SARS-CoV Frankfurt | A | 2 | 150-180 | 12 | mrc/tif 16 bit | 0.96 / 1.17 |
| 5 | SARS-CoV-2 INMI-Italy | B | 3 | 150-180 | 17 | mrc/tif 16 bit | 0.96 / 1.17 |
| 6 | SARS-CoV Frankfurt | A | 5 | 45 | 111 | tif, 16 bit | 0.54 |
| 7 | SARS-CoV-2 INMI-Italy | B | 5 | 45 | 134 | tif, 16 bit | 0.54 |
| 8 | SARS-CoV-2 INMI-Italy | B | 3 | 65 | 85 | tif, 16 bit | 0.64 |
| 9 | SARS-CoV-2 INMI-Italy | B | 3 | 85 | 101 | tif, 16 bit | 0.64 |
| 10 | SARS-CoV-2 INMI-Italy | B | 3 | 110 | 66 | tif, 16 bit | 0.64 |
| 11 | SARS-CoV Frankfurt | A | 4 | 200-250 | 10 | tif, 8 bit | 0.57 |
| 12 | SARS-CoV-2 INMI-Italy | B | 4 | 200-250 | 11 | tif, 8 bit | 0.57 |

The maximal length of the spikes associated with a virus particle was measured by a step-wise (nanometer) extension of the polygonal selection, which was made for each virus particle to determine its maximal diameter. The precision of the method was validated by individual line measurements of spike length.

Extracellular virus particles in thicker (150-180 nm) plastic sections were recorded by single-axis electron tomography using the Tecnai and the bottom-mounted Eagle 4k CCD camera, at a magnification of 18,500x and 23,000x (1.17 and 0.96 nm pixel size) and a binning of 2. Virus particles were selected randomly. If particles appeared morphologically intact and tilting to at least −60 and +60° was possible, a tilt series of the region of interest was recorded. Two datasets, one for SARS-CoV and one for SARS-CoV-2, with a minimum of 12 tilt series each, were recorded (Table 1). Tomograms were reconstructed according to the workflow listed in Supplementary Table 2. Measurements were performed with the Fiji software by using the following workflow. Tomograms were loaded, size calibrated and inspected in the orthoslice view (z, x/z and y/z view). For size measurements, particles were selected which appeared intact, showed no distinct compression by other structures and which were with more than half of their size enclosed in the tomogram volume. Maximal diameter of the selected virus particle (without spikes) was measured by adjusting the z view to a level where the particle in x/z and y/z view becomes maximal in width and by using the oval selection tool with the measurement setting „fit ellipse“. The maximal diameter of the oval (elliptical) selection was noted.

Shrinkage of virus particles in x/y direction during irradiation with the electron beam was measured with the Tecnai using similar electron dose as applied for electron-tomography. Suitable sample positions were selected at low magnification and focusing was done at a distant position (10 μm) using the low-dose module of the Tecnai. Immediately after switching automatically back to target position, a high-resolution image (2048 x 2048 pixel, Eagle camera) was recorded (t = 0 min). The selected sample position was continuously illuminated by the electron beam for 30 min before another image was recorded (t = 30 min). The maximal diameter of individual virus particles was measured at both time points using the “polygonal selection” tool and the “fit ellipse” measurement setting of Fiji.

### Measurement of spike density

The spike number of virus particles was estimated using ultrathin plastic sections of 45 nm thickness, which was the lowest sectioning thickness set point producing regular sections. Extracellular virus particles were randomly selected and recorded with the side-mounted CCD camera at a magnification of 135,000x if the particles met the following criteria: (1) the particle was morphologically intact; (2) the particle was not deformed (e.g. by pressing against other structures); (3) the particle membrane was visible (at least 90% of the perimeter). Two datasets, each with about 150 particles, were recorded (see Table 1). The number of spikes (including partially visible spikes) were counted manually and the maximal diameter of the particles was determined as described in the section before.

Additional measurements were done using complete virus particles extracted from tomograms of 200-250 nm thick sections. We selected all particles which were apparently intact, fully enclosed in the volume of the tomogram and which allowed discrimination of spikes. The volume containing the selected particle was extracted, filtered to increase contrast (Normalize local contrast; maximum pixel size, SD = 5, stretched and centered histogram) and resliced in z to a resolution of 1.5 nm using Fiji. Spikes were labeled and counted manually and the maximal diameter of the virus particles was determined as described in the section before.

## Results

Extracellular virus particles of SARS-CoV and SARS-CoV-2 in Vero cell cultures revealed no significant morphological differences in ultrathin sections (Fig. 2). Virus particles appear as round to oval profiles. A few particles in each population showed another, irregular or deformed particle shape (SARS-CoV: 1.5%, N = 777 particles; SARS-CoV-2: 0.5%, N = 752 particles; Supplementary Fig. S1 A-C). Size distributions of virus particle profiles in conventional ultrathin (65 nm) sections were also similar for both viruses (Fig. 3 A, B). SARS-CoV showed a few larger profiles than SARS-CoV-2, but the medians of the maximal particle diameter without spikes were about the same (SARS-CoV: 95; SARS-CoV-2: 97 nm, without spikes). The replication of the analysis using a second cell culture batch in an independent infection experiment with SARS-CoV-2 resulted in an essentially identical size distribution and median of the maximal particle diameter (Supplementary Fig. S2). We also measured the maximal length of the spikes for each individual virus particle. The median of the measurements was 23 nm for both, SARS-CoV and SARS-CoV-2 (interquartile range of 1 or 2 nm, respectively, N = 100 particles of each virus).

**Figure 2.**
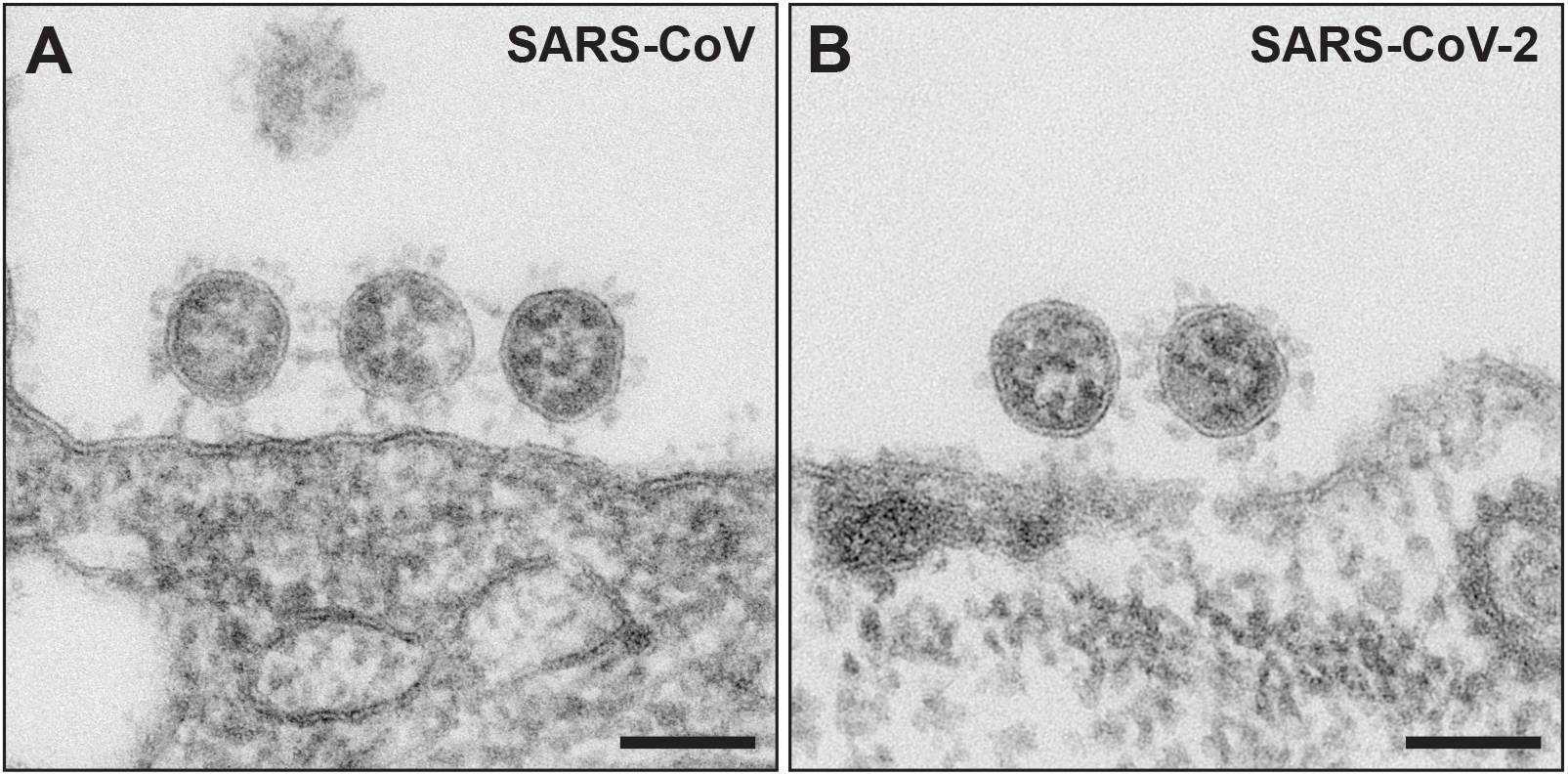
Transmission EM of ultrathin sections through Vero cells which were either infected with SARS-CoV (**A**), or with SARS-CoV-2 (**B**). Viruses are attached to the surface of the cells and do not reveal substantial differences in their ultrastructure. Scale bars = 100 nm.

**Figure 3.**
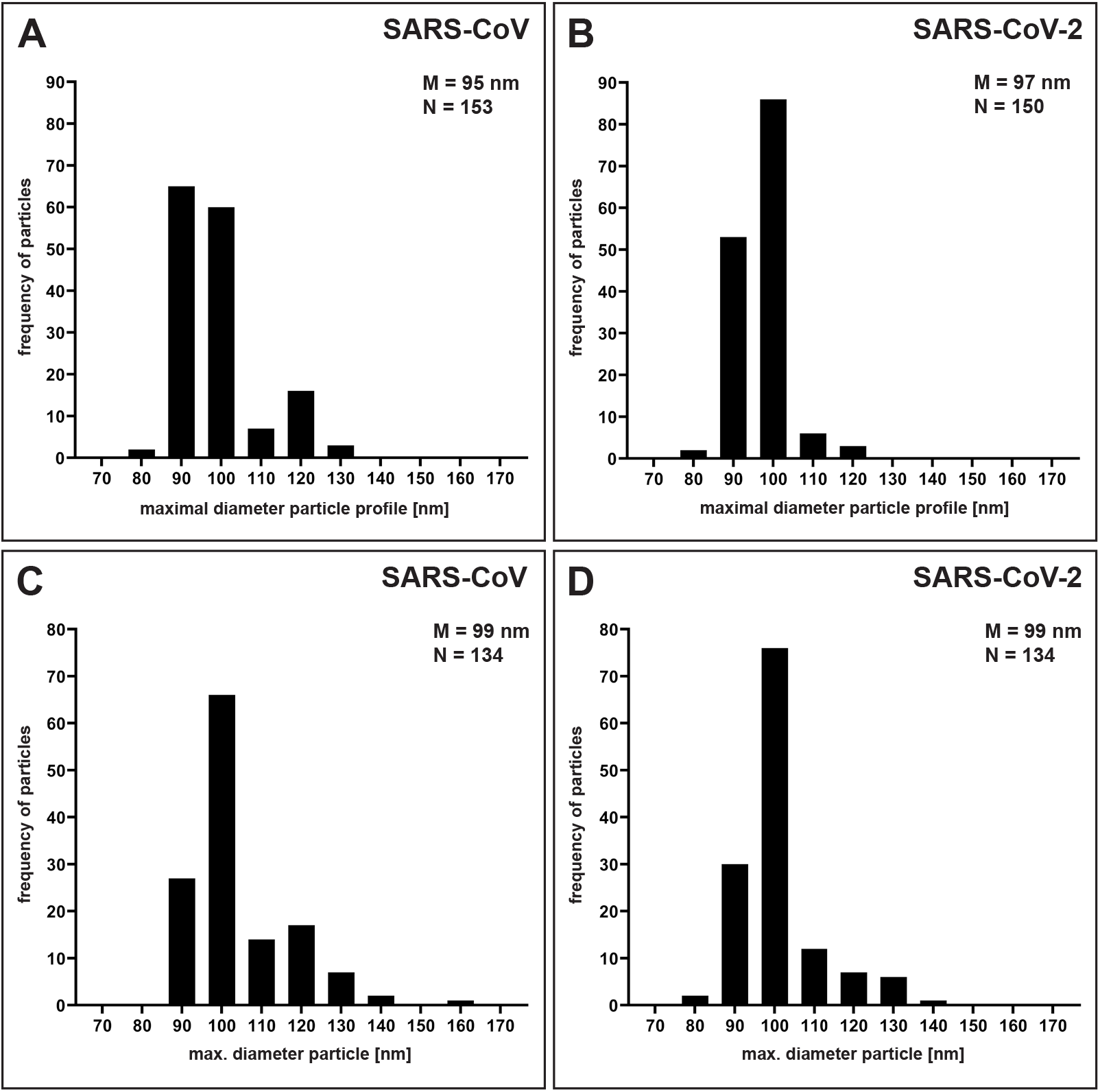
Particle size distribution of SARS-CoV and SARS-CoV-2. **A, B.** Histograms of maximal particle profile diameter without spikes in ultrathin (65 nm) sections (datasets 01 and 02; Table 1). **C, D.** Histograms of maximal particle profile diameter without spikes in electron tomograms of thin (150-180 nm) sections (datasets 04 and 05; Table 1). Particles were measured at their thickest diameter (see Fig. 4 and Methods section). M = median; N = number of measured particles.

The size distribution and the median resulted by measuring the virus particle profiles in conventional ultrathin sections could be biased by an overrepresentation of virus section profiles of a particular virus particle size and/or of undetected deformed particles. Therefore, we recorded tomographic tilt series of viruses in thicker sections (150-180 nm) and calculated single-axis tomograms to measure virus particles at their maximal diameter (Fig. 4) and to exclude deformed (i.e. non-circular/-oval) virus particles. The aligned tilt series and the tomograms showed that almost all of the particles possessed an oval shape (Supplementary Videos 1-4). We rarely detected deformed particles in the tomograms (SARS-CoV: 4%, N = 341 particles; SARS-CoV-2: 2%, N = 276 particles). However, the fraction was slightly higher than measured for particles in sections (see above). In the SARS-CoV samples we found one small cluster of deformed viruses attached to a cell (Supplementary Fig. S1 D) which was excluded from the measurements. The particle size distribution determined in tomograms is similar to the particle size distribution measured in ultrathin sections (Fig. 3 A-C), with an identical median for SARS-CoV and SARS-CoV-2 of 99 nm, which is a few nanometers higher than measured in ultrathin sections of 65 nm thickness. The size distribution shows a slight shift to higher particle diameter for the SARS-CoV (Fig. 3 C, D). We have to note that the thin sections shrunk during electron beam illumination which caused a compressed appearance of the particles in x/z and y/z direction (Fig. 4). This effect is well known and usually does not affect dimensions in x/y if samples/sections are well fixed at their supports^23^, which most likely was the case during our image recording because we used sections on grids with rather small holes and finally stabilized the sections by a carbon layer. This assumption is supported by measurements of virus particle shrinkage in x/y during 30 min of electron beam exposure, which revealed a mean shrinkage of about −0.5 % (SD = 3 %, N = 39 particles, N = 6 exposure experiments) between t = 0 and t = 30 min.

**Figure 4.**
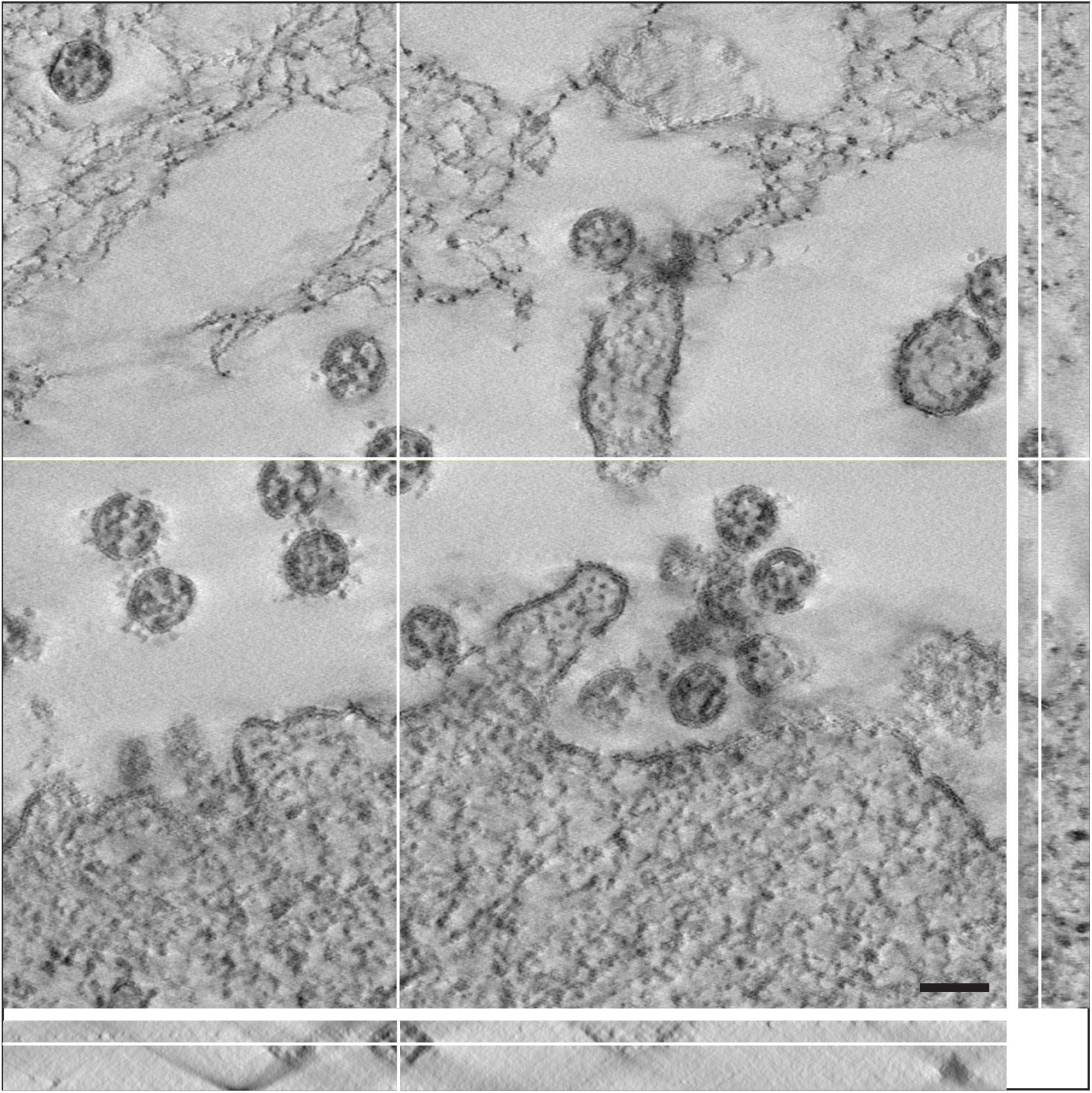
A single digital slice (z view) of an electron tomogram of SARS-CoV-2 particles. The ortho-slice view shows the particle labelled by the white cross lines in side view (x/z and y/z) of the volume at the indicated section plane. The particle appears ovoid in shape and the thickest part of the particle in z was selected for size measurement. Note that the section is compressed in z and thinner than the nominal 180 nm set at the microtome, which also affects the shape of the particle viewed in x/z and y/z. This artifact is well known in electron tomography of plastic sections and only slightly affects the size in x/y^23^ which our shrinkage measurements demonstrated (see Results). Scale bar = 100 nm.

To check whether the different size values measured in section profiles and in tomograms were caused by a variable distortion of the sections at different thickness settings, we measured particle shape descriptors of virus particles in sections of different section thickness. The results showed that the slightly oval particle shape (aspect ratio of about 1.1) is independent of the section thickness in the investigated range of 45 to 110 nm (Table 2), which indicates that the overall shape of the virus particles is rather constant over a wide range of section thickness. The size distribution of the maximal particle diameter at different section thickness is shown in Supplementary Figure S3 and approaches the distribution measured for particles in tomograms with increasing section thickness.

**Table 2.** Shape descriptors of virus particle profiles at different section thickness

| Virus | Data set | Section thickness [nm] | Aspect ratio | Circularity |
| --- | --- | --- | --- | --- |
| SARS-CoV | 06 | 45 | 1.1 | 0.98 |
| SARS-CoV-2 | 07 | 45 | 1.1 | 0.97 |
| SARS-CoV-2 | 08 | 65 | 1.1 | 0.97 |
| SARS-CoV-2 | 09 | 85 | 1.1 | 0.98 |
| SARS-CoV-2 | 10 | 110 | 1.1 | 0.98 |

To get an idea about the spike number on the two different coronaviruses, we counted the spikes present on particle profiles in plastic sections of 45 nm thickness and related their number to the maximal diameter of the individual profile. Figure 5 A and B shows two representative virus particles of the datasets. The scatter plot of the spike number per virus particle diameter for the two coronaviruses revealed a similar shape with an accumulation of size values around 100 nm but a slightly shifted median of the spike number, i.e. SARS-CoV = 12 and SARS-CoV-2 = 10 spikes per particle (Fig. 5 C, D). To get an idea whether this result and measurement approach represent the particle number of entire virus particles, we recorded tomograms of thick (200-250 nm) plastic sections and counted the spikes of complete virus particles. The results revealed a similar tendency than the measurements of the spike number associated with virus section profiles (Fig. 5 C-F), with a higher spike number for SARS-CoV (M = 32 spikes per virus particle) than for SARS-CoV-2 (M = 25 spikes per virus particle). Although the distributions of the spike number were widely overlapping (Fig. 5 C-F), measurements indicated that the investigated SARS-CoV virus population carried more spikes at their surface than the SARS-CoV-2 virus population.

**Figure 5.**
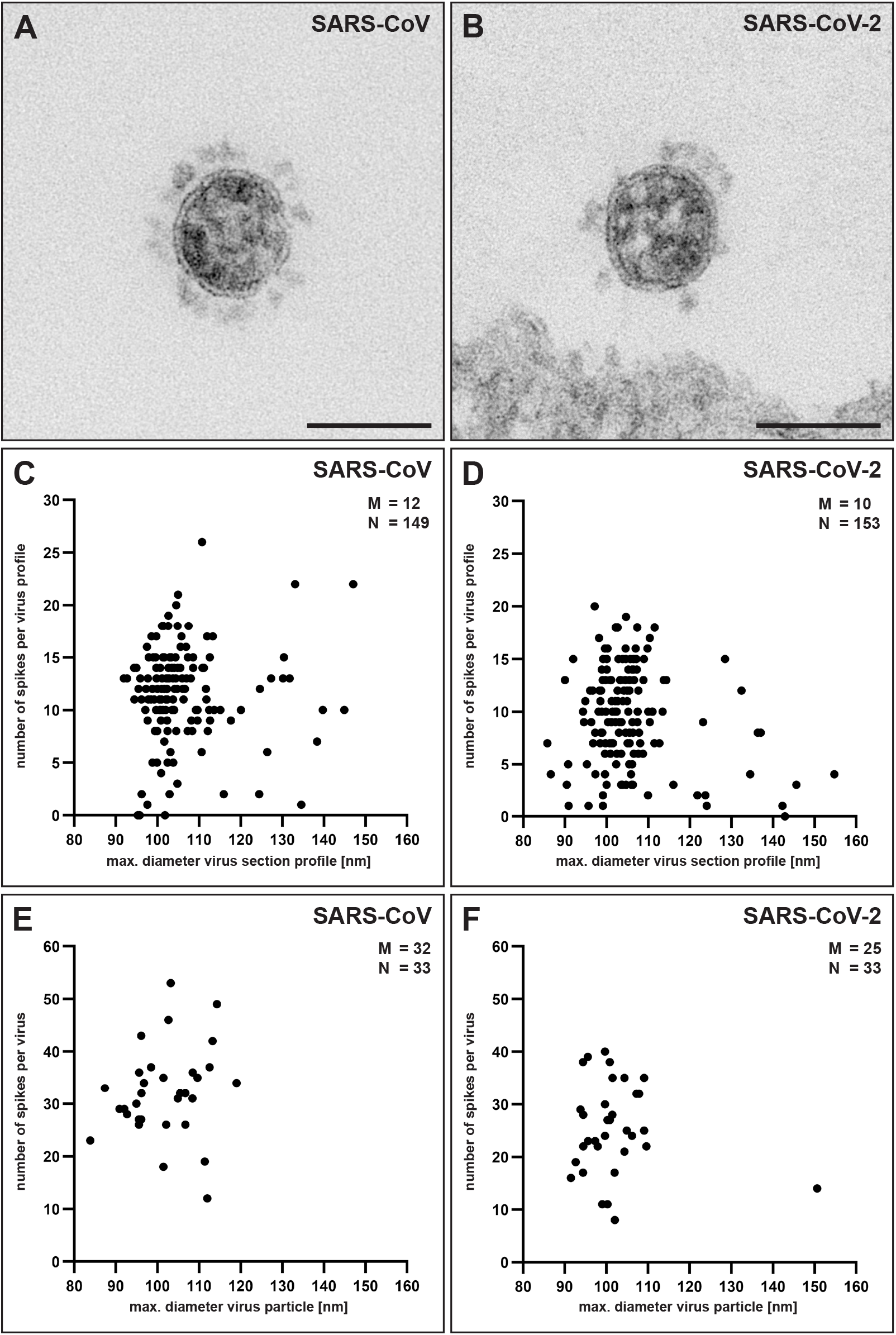
Determination of the spike number of SARS-CoV and SARS-CoV-2 by transmission EM. **A,B.** Single virus particles of either SARS-CoV (**A**) or SARS-CoV-2 (**B**) in ultrathin (45 nm) sections which show differences in their spike number. Scale bars = 100 nm. **C, D.** Scatter plot of the number of spikes per maximal particle profile diameter of SARS-CoV (**C**) and SARS-CoV-2 (**D**) (datasets 06 and 07; Table 1) in ultrathin (45 nm) sections. **E, F.** Scatter plot of the number of spikes per maximal particle diameter of SARS-CoV (**E**) and SARS-CoV-2 (**F**) (datasets 11 and 12; Table 1) in tomograms of thin (200-250 nm) sections. M = median; N = number of measured particles.

## Discussion

We determined the size distribution and the spike number of SARS-CoV and SARS-CoV-2 virus particles *in situ,* in the surrounding of virus producing Vero cells, by using thin section EM. Viruses and cells were chemically inactivated and stabilized by glutaraldehyde *in situ* and embedded in plastic. This preparation procedure changes the ultrastructure of biological objects^24^, including their dimensions^25^, e.g. by adding chemicals or by removing the water, and it does not preserve their accurate molecular structure^26^. However, at the resolution level sufficient to study the ultrastructure of organelles (i.e. their shape and internal architecture), this procedure provides reliable information which is, at this resolution, in many cases very similar to the information obtained by cryo-EM^24^, the gold standard in structural biology.

Cryo-EM provides maximal structural information about the virus architecture down to the molecular level^27,28^. However, for single particle cryo-EM, virus particles usually have to be concentrated and purified, which is not trivial, especially for enveloped viruses. Purification and/or enrichment can select for a certain particle size and shape, introduce deformations^29^, which was also observed for SARS-CoV-2^11^, and might cause loss of membrane protein^30^. Biosafety still requires inactivation of the virus preparation before conducting the sample preparation for cryo-EM, and the effects on the ultrastructure must be carefully controlled. The recently published work on isolated SARS-CoV-2^11–13^ only partially addressed those aspects^11^.

Studying virus particles by cryo-EM *in situ* attached to or present in the cells is extremely difficult to perform, since whole cell cryo-EM (i.e. cryo-electron tomography) needs either thin parts of an infected cell or lamella preparation by FIB-SEM to generate datasets of frozen hydrated and therefore virtually unchanged virus particles^31^. This work is technically extremely challenging and very time consuming^31^ and usually restricted to a limited set of samples which not necessarily fully represent the biological variability of the sample. However, in a recent study both approaches could be applied to SARS-CoV-2 infections of different cell lines^14^ providing valuable structural data on cell-associated virus.

Our study was intended to provide a reference for ultrastructural work performed on virus infected cells embedded in plastic, because this method is widely used to study, for instance, the cell biology of infection models or infected patient material. The results revealed that SARS-CoV and SARS-CoV-2 are very similar in morphology and size, as could be expected from the close taxonomic relationship of the two viruses^4^ and reports on the virus ultrastructure in plastic sections which are available^8,9,15^. However, the similarity of the size distribution of the two coronaviruses tested and of the two biological replicates (two independent infection experiments with SARS-CoV-2) was a surprise because enveloped viruses are usually more variable in shape and size than non-enveloped viruses^32^. The median maximal diameter of the virus particles which we determined in sections is in the wide range of particle sizes reported in the literature (60-140 nm)^33–36^. The large variability of the reports could be due to differences in the measurement techniques used or to variations of the ultrastructural preservation achieved by the various fixation and embedding protocols. The tannic acid and uranyl acetate *en bloc* contrasting applied in our preparations may have increased the particles artificially. However, preliminary experiments indicate that the effect on the measurement of the particle size is small (Supplementary Table 3) and most probably caused by a reduced visibility of the virus particle membrane in samples which were only treated by osmium tetroxide.

We used two different strategies for determination of virus particle size in thin sections: (1) Measurement of virus particle section profiles in ultrathin (65 nm) sections and (2) measurement of widest particle profile in tomograms of thin (150-180 nm) sections. The size distribution median was a few nanometers higher in tomograms (99 nm) than in ultrathin sections (95 and 97 nm). Since shape descriptors indicate a constant particle shape in thin and thicker ultrathin sections and measurement of shrinkage in x/y-direction during long-time irradiation showed no significant shrinkage of virus particles during recording of tomography tilt series, we conclude that the particle size measured in tomograms represent the correct size of virus particles in plastic sections. Median and shape of the size distribution of particles in ultrathin sections and of particles in tomograms of thicker sections converge with increasing thickness of ultrathin section. The difference between the two measurement approaches can be explained by an over-and/or underrepresentation of particle section profiles of a particular size class at a particular section thickness. However, our results show that measurement of particle size in ultrathin sections of the standard section thickness between 65 and 110 nm provides a good estimator for the size of the coronavirus particles embedded in plastic.

The size values measured for SARS-CoV in our study (~100 nm, without spikes) differ from the values measured by cryo-EM (SARS-CoV: 82-94^37,38^; SARS-CoV-2: 90-97 nm^11,13,14^). As already mentioned above, it is highly likely that the plastic embedding changed the shape and size of the virus particles. Obviously, the virus particles in thin plastic sections appear more oval than the particles shown by cryo-EM^11,12,14^. Therefore, a simple explanation for the difference of the particle size measured by thin section EM and cryo-EM could be the change of particle shape from round to oval in thin section EM. The aspect ratio of the virus particle enveloping ellipse in thin section EM was about 1.1 over a wide range of section thickness which indicates an oval particle shape. A change from round (aspect ratio = 1) to an oval shape with the aspect ration 1.1 would be associated with an increase in maximal particle diameter of about 10% which roughly amounts the difference between particle size measured by cryo-EM (~90 nm) and thin section EM (~100 nm). Other reasons to explain the difference could be the different virus strains which we have used in comparison to the strains used in the cryo-EM studies^11,13,14,37,38^ or differences in the cell culture which seem to have an effect on the size distribution^11^. It is also not possible to exclude that concentration and purification of virus particles before cryofixation have an impact on the size distribution of the virus particle population. A comparison of non-purified and purified SARS-CoV-2 showed only small differences (91 vs. 92 nm^11^) and the measurement of SARS-CoV-2 *in situ* resulted in similar values (90 nm^14^) which suggest that virus particles were not affected during preparation in those experiments.

The maximal length of the spikes associated with the virus particles in ultrathin sections was 23 nm in SARS-CoV and SARS-CoV-2, which is close to the values determined by Cryo-EM (SARS-CoV: 19 nm^38^, SARS-CoV-2: 25 nm^14^). However, it is possible that the tannic acid, which was used for *en bloc* contrasting, has increased the spike size artificially because tannic acid is known to bind to glycoproteins^39^.

The measurement of the spike number associated with virus particle profiles in ultrathin sections revealed differences between the SARS-CoV and SARS-CoV-2 virus populations studied, which could be supported in their tendency by determination of the spike number of entire virus particles in tomograms. A qualitative difference of the spike density between SARS-CoV and SARS-CoV-2 was already observed by the study of Ogando *et al.*^9^ and associated with a reduced infectivity of SARS-CoV-2 in comparison to SARS-CoV. Our quantitative measurements, which were performed with the same SARS-CoV isolate but a different SARS-CoV-2 isolate than the one used in the study of Ogando *et al.*^9^, support this conclusion. For SARS-CoV, Beniac *et al.*^37^ estimated a mean number of 65 spikes per virus using cryo-EM, with a certain variability in distribution between different particles, which is much higher than the median number which we have measured. Again, Beniac *et al.*^37^ used a different SARS-CoV isolate than we have used in our study (Tor 3 versus Frankfurt 1), which may explain the observed differences. Spike density of SARS-CoV-2 was determined recently by cryo-EM for different virus isolates than the virus isolate used in our study. The reported mean values vary from 25 to 40 spikes per virus particle with significant variation among particles^11–14^. We also detected a high variability of the spike number among virus particles and with a median of 25 spikes per virus particle our measurements fit to the lower values measured for the other isolates.

The relevance of the observed difference in spike number between different virus isolates is not known but could be related to virus infectivity and fitness. Our results show that these differences can be detected by measuring the spike number in thin sections, which is much easier than by using (cryo) electron tomography. Further studies should measure the spike number of isolates already present or rapidly evolving in the human population^40^ and relate it with virus infectivity and receptor-binding affinity, to get an idea if adaptation of the spike protein is responsible for the fitness of particular virus isolate in the population.

In summary, we provide morphometric data for SARS-CoV and SARS-CoV-2 particles in plastic sections, which are very similar to the data obtained by cryo-EM. All raw datasets can be used for re-investigation or other purposes (e.g. for validation / testing / training of computer algorithms). The major outcome is that the investigated isolates of SARS-CoV and SARS-CoV-2 are ultrastructurally very similar in shape and size and show a small difference in their spike number.

## Data availability

Datasets 01 to 12 (see Table 1) are available at the data repository Zenodo:

Dataset 01: DOI 10.5281/zenodo.3985098
Dataset 02: DOI 10.5281/zenodo.3985103
Dataset 03: DOI 10.5281/zenodo.3985110
Dataset 04: DOI 10.5281/zenodo.3985120
Dataset 05: DOI 10.5281/zenodo.3985424
Dataset 06: DOI 10.5281/zenodo.3986526
Dataset 07: DOI 10.5281/zenodo.3986580
Dataset 08: DOI 10.5281/zenodo.4275703
Dataset 09: DOI 10.5281/zenodo.4275728
Dataset 10: DOI 10.5281/zenodo.4275735
Dataset 11: DOI 10.5281/zenodo.4275742
Dataset 12: DOI 10.5281/zenodo.4275769

## Acknowledgements

We would like to thank Silvie Muschter and Annette Teichman for conducting the cell culture and Gudrun Holland, Petra Kaiser and Freya Kaulbars for embedding of the samples. We are also grateful to Christoph Schaudinn for reading of the manuscript and his valuable suggestions and to Ursula Erikli for copy-editing. Finally, the authors would particularly acknowledge the continuous efforts of Hans Gelderblom in the past to improve ultrastructural analysis of viruses in our laboratory and beyond.

## Author contributions

M.L. designed the study and wrote the manuscript; A.K., T.H., L.M. and M.L. performed the EM investigations; J.M. and A.N. planned the cell culture and infection experiments, including their quality assurance. All authors discussed the results and their presentation and approved the final version.

## Additional information

Supplementary information accompanies this paper.

All image data used for the measurements are available at the Zenodo research data repository.

