## Supplementary Information for "Morphometry of SARS-CoV and SARS-CoV-2 particles in ultrathin plastic sections of infected Vero cell cultures"

### Supplementary Methods

**Supplementary Table S1.** Stepwise embedding protocol for thin section EM (Laue 2010)

| Step | Solution/Mixture/Solvent | Duration of incubation [min] | Temperature | Tissue processor |
|---|---|---|---|---|
| 1 | Hepes buffer, 0.05 M | 5 | RT | no |
| 2 | Hepes buffer, 0.05 M | 5 | RT | no |
| 3 | Hepes buffer, 0.05 M | 5 | RT | no |
| 4 | Osmium tetroxide, 1% in water | 60 | RT | no |
| 5 | Distilled water | 5 | RT | no |
| 6 | Distilled water | 5 | RT | no |
| 7 | Distilled water | 5 | RT | no |
| 8 | Tannic acid, 0.1% in 0.05 M Hepes buffer | 30 | RT | yes |
| 9 | $Na_2SO_4$ 1% in 0.05 M Hepes buffer | 10 | RT | yes |
| 10 | $Na_2SO_4$ 1% in 0.05 M Hepes buffer | 10 | RT | yes |
| 11 | Distilled water | 10 | RT | yes |
| 12 | Distilled water | 10 | RT | yes |
| 13 | Distilled water | 10 | RT | yes |
| 14 | Uranyl acetate, 2% in distilled water | 120 | RT | yes |
| 15 | Ethanol, 30% | 30 | RT | yes |
| 16 | Ethanol, 50% | 30 | RT | yes |
| 17 | Ethanol, 70% | 30 | RT | yes |
| 18 | Ethanol, 95% | 60 | RT | yes |
| 19 | Ethanol, abs. | 60 | RT | yes |
| 20 | Ethanol, abs. | 60 | RT | yes |
| 21 | Propylene oxide | 30 | RT | yes |

| | | | | |
|---|---|---|---|---|
| 22 | Propylene oxide | 30 | RT | yes |
| 23 | Propylene oxide / Epon 2+1 | 180 | RT | yes |
| 24 | Propylene oxide / Epon 1+1 | 180 | RT | yes |
| 25 | Propylene oxide / Epon 1+3 | 180 | RT | yes |
| 26 | Epon | 120 | RT | yes |
| 27 | Epon | 180 | RT | yes |
| 28 | Epon | 240 | RT | no |
| 29 | Epon - final embedding in silicone moulds | - | RT | no |
| 30 | Polymerization | 2 days | 60 °C | no |

Tissue processor = Leica EM TP (Leica Microsystems)

**Supplementary Table S2.** Workflow and parameters for alignment and reconstruction of electron tomograms using Inspect3D

| Step | Inspect3D Tasks | Parameter | Value |
|---|---|---|---|
| 1 | Reduce data | Removal of hot pixels/Removal Pass | 1 |
| | | Removal of hot pixels/slider | 800 |
| | | Remove unsharp/mis-exposed images | Variable (at least 120 images should remain) |
| 2 | Setup filter | High pass | 0.02 |
| | | Radius | 0 |
| | | Low Pass | 0.6 |
| | | Radius | 0 |
| | | Tapering | 0 |
| | | Add Hanning window | Yes |
| 3 | Calculate alignment shifts | Repeat until pixel shifts approach zero | Variable (5-6 repetitions) |
| | | Use filter | Yes |
| | | Use stretching | Yes |
| | | Add measurement to current shifts | Yes (after first run) |
| 4 | Apply alignments | Use shifts from alignment data view | Yes |

| | | | |
|---|---|---|---|
| | | Use corrections from tilt axis alignment task | No |
| 5 | Apply alignments | Apply suggested additional correction of tilt axis | Variable (according to tilt axis determination by the FFT) |
| 6 | Tilt axis adjustment | Rebin Input Size | 512 |
| | | Stack | miP |
| | | Shift and tilt | variable (Reduction of "banana"-shaped structures in reconstructed sections) |
| 7 | Apply alignments (select last aligned file as input) | Use shifts from alignment data view | No |
| | | Use corrections from tilt axis alignment task | Yes |
| 8 | Reconstruction | SIRT | Iterations = 25 |

**Supplementary Table S3.** Maximum virus particle diameter of SARS-CoV-2 in samples processed with different *en bloc* contrasting procedures (medians of 50 particles each)

| ***en bloc* contrasting** | **Maximal particle diameter [nm]** |
|---|---|
| Os Ta UA | 97 |
| Os UA | 99 |
| Os | 94 |

Os = Osmium tetroxide
Ta = Tannic acid
UA = Uranyl acetate

**Thin section electron microscopy with or without *en bloc* contrasting**

Processing of the cells was the same as described in the Methods section with the exception that the samples were processed manually and not in a tissue processor. Incubation time for dehydration and resin infiltration were shortened, but not for the post-fixation with osmium tetroxide and the *en bloc* contrasting with tannic acid and uranyl acetate. Three variants were prepared from cells which were used to produce sample B (Table 1): (1) osmium tetroxide, tannic acid and uranyl acetate (reference sample); (2) osmium tetroxide, uranyl acetate; (3) osmium tetroxide. Ultrathin sections (65 m) were produced and treated as described in the Methods section. Transmission electron microscopy was performed with the Tecnai and the Megaview camera. Measurements of the maximal particle profile diameter were conducted as described in the Methods section.

### Supplementary Figures

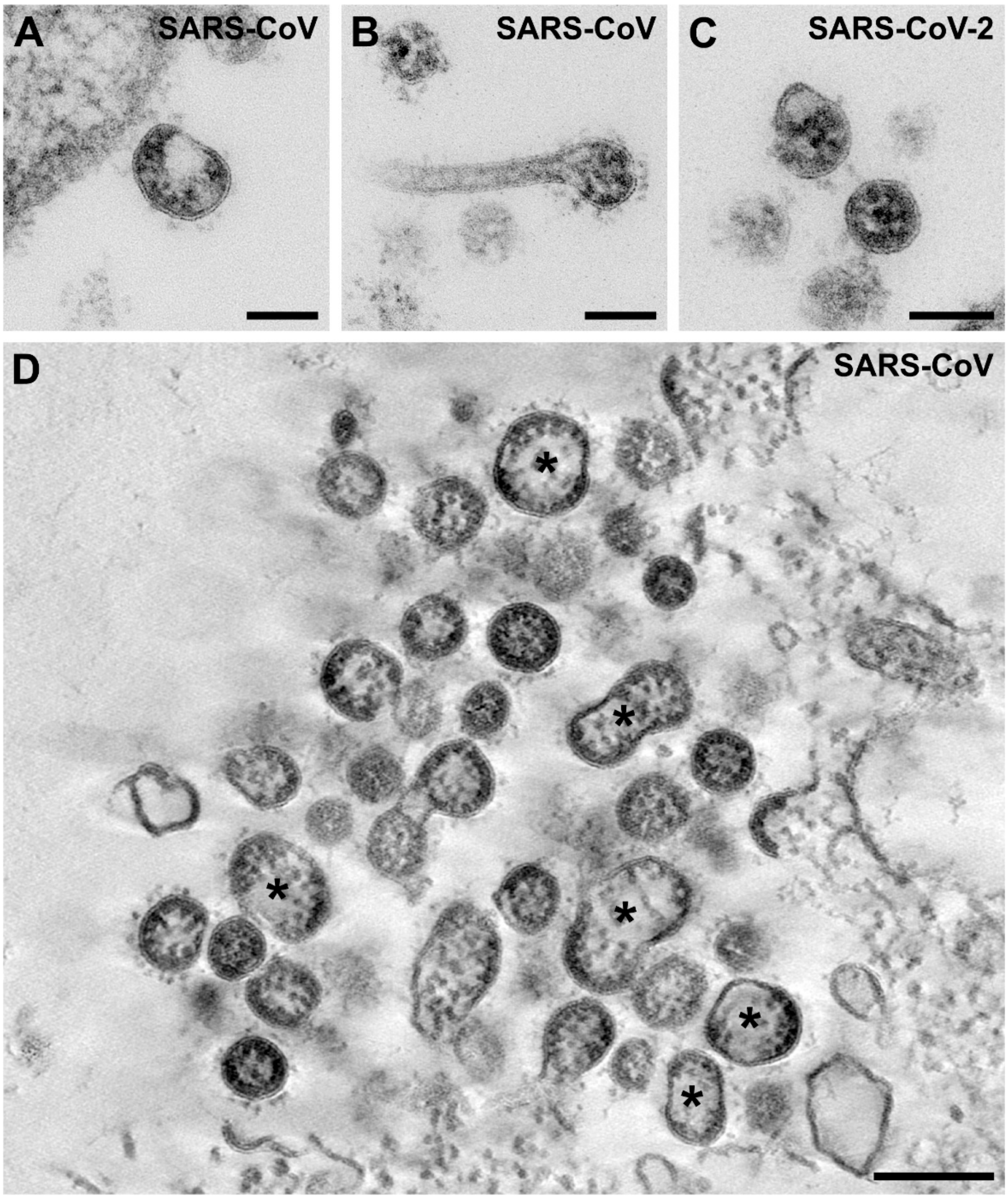


**Supplementary Figure S1.** Deformed virus particles of SARS-CoV (**A, B, D**) and SARS-CoV-2 (**C**), which were excluded from the particle size analysis (see Results section for more details on the frequency of deformed particles). **A-C** Deformed virus particles in ultrathin sections. **D** A higher number of deformed virus particles (*) at the surface of a Vero cell. Single slice of a tomogram of a thin plastic section. Scale bar in A-C = 100 nm and in D = 200 nm.

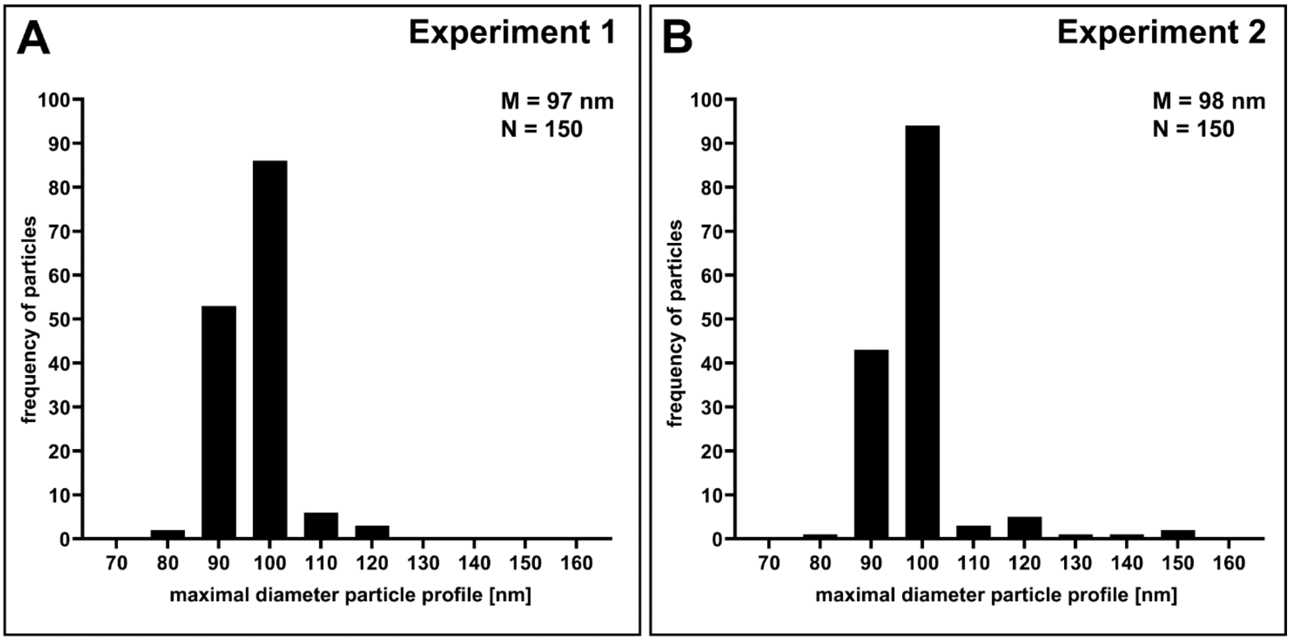


**Supplementary Figure S2.** Comparison of virus particle size distributions of two virus populations generated by two independent cell infection experiments (**A**, Experiment 1**; B**, Experiment 2) using SARS-CoV-2 (Italy-INMI1) and Vero cells. The maximal diameter of particle profiles without spikes was measured in ultrathin sections (datasets 02 and 03; Table 1). M = median; N = number of particles measured.

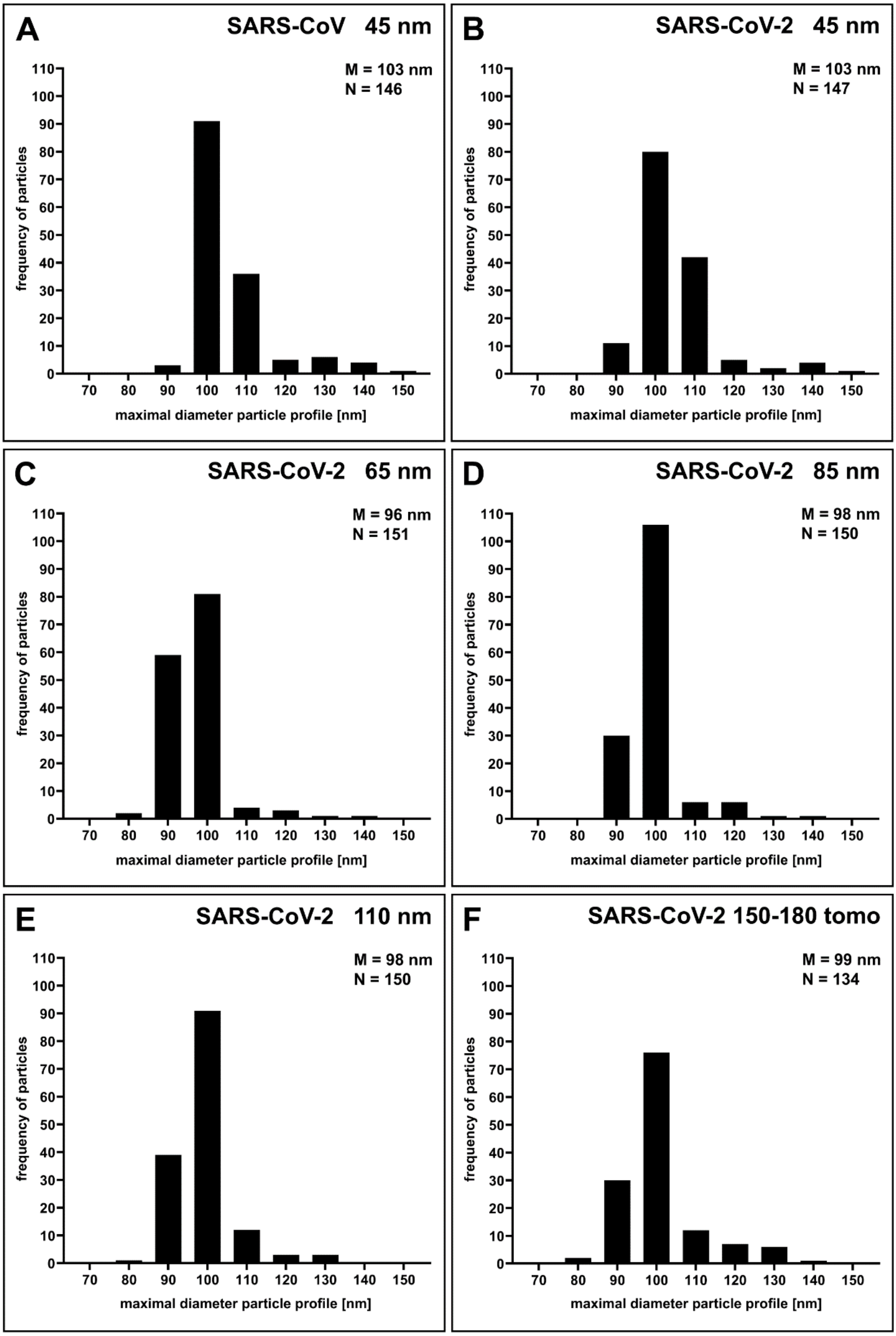


**Supplementary Figure S3.** Size distribution of SARS-CoV and SARS-CoV-2 particle profiles in ultrathin sections of different section thickness. The maximum diameter of the particle profile was measured. **A**. 45 nm thick sections of SARS-CoV (dataset 06, Table 1). **B**. 45 nm thick sections of SARS-CoV-2 (dataset 07, Table 1). **C**. 65 nm thick sections of SARS-CoV-2 (dataset 08, Table 1). **D**. 85 nm thick sections of SARS-CoV-2 (dataset 09, Table 1). **E**. 110 nm thick sections of SARS-CoV-2 (dataset 10, Table 1). **F**. 150-180 nm thick sections of SARS-CoV-2 (dataset 05, Table 1; same data as shown in Figure 3 D, but displayed at different range).

### Supplementary Videos

**Supplementary Video 1.** Aligned image tilt series of a thin section through SARS-CoV particles at the surface of a Vero cell. The corresponding digital sections through the calculated tomogram are shown in the Supplementary Video 2. Use the loop function of your video player for continuous replay.

**Supplementary Video 2.** Digital sections (z = 1.17 nm) through a tomogram calculated from the image tilt series shown in the Supplementary Video 1. Use the loop function of your video player for continuous replay.

**Supplementary Video 3.** Aligned image tilt series of a thin section through SARS-CoV-2 particles at the surface of a Vero cell. The corresponding digital sections through the calculated tomogram are shown in the Supplementary Video 2. Use the loop function of your video player for continuous replay.

**Supplementary Video 4.** Digital sections (z = 1.17 nm) through a tomogram calculated from the image tilt series shown in the Supplementary Video 3. Use the loop function of your video player for continuous replay.
